# DARQ: Deep learning of quality control for stereotaxic registration of human brain MRI

**DOI:** 10.1101/2021.08.16.456514

**Authors:** Vladimir S. Fonov, Mahsa Dadar, The PREVENT-AD Research Group, ADNI, D. Louis Collins

## Abstract

Linear registration to stereotaxic space is a common first step in many automated image-processing tools for analysis of human brain MRI scans. This step is crucial for the success of the subsequent image-processing steps. Several well-established algorithms are commonly used in the field of neuroimaging for this task, but none have a 100% success rate. Manual assessment of the registration is commonly used as part of quality control. To reduce the burden of this time-consuming step, we propose Deep Automated Registration Qc (DARQ), a fully automatic quality control method based on deep learning that can replace the human rater and accurately perform quality control assessment for stereotaxic registration of T1w brain scans.

In a recently published study from our group comparing linear registration methods, we used a database of 9325 MRI scans and 64476 registrations from several publicly available datasets and applied seven linear registration tools to them. In this study, the resulting images that were assessed and labeled by a human rater are used to train a deep neural network to detect cases when registration failed. We further validated the results on an independent dataset of patients with multiple sclerosis, with manual QC labels available (n=1200).

In terms of agreement with a manual rater, our automated QC method was able to achieve 89%accuracy and 85% true negative rate (equivalently 15% false positive rate) in detecting scans that should pass quality control in a balanced cross-validation experiments, and 96.1% accuracy and 95.5% true negative rate (or 4.5% FPR) when evaluated in a balanced independent sample, similar to manual QC rater (test-retest accuracy of 93%).

The results show that DARQ is robust, fast, accurate, and generalizable in detecting failure in linear stereotaxic registrations and can substantially reduce QC time (by a factor of 20 or more) when processing large datasets.

## 1 Introduction

Many automatic image-processing techniques to analyze human brain MRI scans include linear registration to stereotaxic space as one of the first steps in the pipeline (Fischl, 2012; Jenkinson et al., 2012; Zijdenbos et al., 2002). Successive image processing steps are highly dependent on the success of the stereotaxic registration. Often, a human expert rater must manually verify the quality of this step by looking at a series of images that show overlap between the registered scan and a reference template to identify datasets that failed registration. Such failed datasets may be re-registered with different parameter settings, be subject to subsequent manual registration, or be discarded from analysis. As explicitly stated by Ashburner et al. (Ashburner and Friston, 2000), registration quality should be maintained as high as possible. Depending on the dataset and registration algorithm used, the success rate can vary from almost 100% down to 60% (Dadar et al., 2018). In particular, MRI scans of subjects with space occupying lesions or those with strong atrophy due to neurodegenerative diseases show lower rates of success when using registration tools that were originally developed and tested on MRI scans of young healthy subjects (Dadar et al., 2018). Also, our experiments show that strong atrophy makes the task of manual quality control even more challenging when the reference template is representative of a healthy population.

In a previous paper from our group comparing reliability of linear registration algorithms (Dadar et al., 2018), we performed an experiment where a large number of scans (9325 scans, 3308 unique subjects) from the Human Connectome Project (HCP) (Van Essen et al., 2012), the Alzheimer’s Disease Neuroimaging Initiative (ADNI) (Mueller et al., 2005), the Pre-symptomatic evaluation of experimental or novel treatments for Alzheimer’s Disease (PreventAD), and the Parkinson’s Progression Marker Initiative (PPMI) (Tremblay-Mercier et al., 2014) databases were linearly registered to the MNI-ICBM152 2009c space (Fonov et al., 2011; Manera et al., 2020) using five publicly available linear registration methods. All registrations were then assessed for quality by a human rater. Since manual quality control of registrations is time consuming (~30 hours for 9693 registrations) and prone to interrater and intra-rater errors (e.g., QC intra-rater Dice overlap index of 0.96 (Dadar et al., 2018)), an automatic method that would be able to assess quality of registrations accurately and consistently would be useful for the community.

Given the emergence of large MRI datasets such as the UKBioBank (UKBB), manual inspection of registration quality becomes unfeasible. Recently, several papers have been published proposing methods for automatic or semi-automatic quality control in large studies (Alfaro-Almagro et al., 2018; de Senneville et al., 2020; Dubost et al., 2020). Of particular interest is the work of Alfaro-Almago et al., where a single pass/fail metric is used to summarize the quality of the MRI acquisition and preprocessing (Alfaro-Almagro et al., 2018). This method was applied to the UKBB project where the number of acquired scans is expected to be more than 100,000 (Sudlow et al., 2015). In contrast to this approach, we propose a method focusing on the quality control of a single important step used in many preprocessing pipelines: automatic linear registration of T1w MRI scans into stereotaxic space. Recently, de Senneville et al. proposed an automated registration QC method known as RegQCNET to solve the same problem that uses a deep learning-based estimate of misregistration distance trained on simulated misregistrations (de Senneville et al., 2020). A threshold on the estimated distance can be used to pass or fail datasets. The main difference of our approach is that we use a large database of MRI scans that were manually inspected by an experienced rater, with registration mistakes corresponding to those produced by several widely used registration tools whereas the method by Senneville et al. uses artificially generated data.

Another recently published paper by Dubost et al. (Dubost et al., 2020) addresses the problem in a different fashion; by calculating an overlap metric between segmentation masks of anatomical features (i.e., the lateral ventricles) and the anatomical atlas after registration. This method uses a different imaging modality (FLAIR) and relies on the automatic segmentation of anatomical features without registration. In our opinion, this approach relies on an additional unnecessary step (segmentation of lateral ventricles without linear registration) that could be another point of error or failure. In somewhat similar QC applications, Benhalaji et al. evaluated both manual expert and crowdsourcing techniques for stereotaxic registration QC, but have not developed an automatic QC method (Benhajali et al., 2020). Other QC systems exist, but they are not applied to stereotaxic registration. For example, Küstner et al have developed a reference-free method to evaluate MRI acquisition quality using deep learning (Küstner et al., 2018) and the LONI QC system produces a number of QC metrics that can be used to automatically evaluate MRI acquisitions (Kim et al., 2019).

Our goal was to design, build, and test a system to automatically identify datasets that pass or fail stereotaxic registration QC with very high certainty (here we operationally define certainty as True Negative Rate) so that manual QC can focus only on a much smaller subset of questionable cases. Given that many registration algorithms have a greater than 90-95% success rate (Dadar et al., 2018), such a QC tool could reduce the manual QC effort by a factor of 10 to 20, a significant savings of time and energy. We attempted to replicate the behavior of a human rater in this task by training a deep neural network (DNN) to determine quality of registrations by analyzing a series of 2D control images. Following the logic from (Ashburner and Friston, 2000), we determined that it is more important to ensure the quality of the images that were accepted by the automatic rating tool, rather then maximize the overall accuracy of agreement with the manual classification of pass/fail. We thus decided to minimize the False Positive Rate (FPR) metric (reflecting the proportion of incorrect registrations that were passed by the automated QC tool) or equivalently maximizing the True Negative Tate (TNR). In order to leverage existing network design, and to speed training of the DNN, we adapted a pre-trained network that behaved well on image classification tasks (Canziani et al., 2016), the Resnet-18 (Gross and Wilber, 2016; He et al., 2016).

In our paper, we also compared our classification approach to the misregistration estimation approach by de Senneville et al. (de Senneville et al., 2020). Based on our experimental results, misregistration distance estimation achieved lower performance than direct classification. Finally, to allow others to take advantage of our proposed automated QC tool, we have made DARQ publicly available at https://github.com/vfonov/DARQ

## 2 Methods and materials

### 2.1 Materials

We used T1w MRI scans from four different datasets (9693 scans in total):

- **ADNI:** The Alzheimer’s Disease Neuroimaging Initiative (ADNI) (Mueller et al., 2005), is a multi-center and multi-scanner study with the aim of defining the progression of Alzheimer’s disease (AD). Subjects are normal controls, individuals with mild cognitive impairment (MCI) or AD aged 55 years or older. Data was acquired using 1.5T and 3T scanners of different models of GE Medical Systems, Philips Medical systems, and SIEMENS at 59 acquisition sites. We used ADNI1, ADNI2, and ADNI/GO data, including 3136 T1-weighted scans from 1.5T scanners and 3041 scans from 3.0T.
- **PPMI**: The Parkinson Progression Marker Initiative (PPMI) (Marek et al., 2011) is an observational, multi-center and multi-scanner longitudinal study designed to identify PD biomarkers. Subjects are normal controls or de Novo Parkinson’s patients aged 30 years or older. Data was acquired using 1.5T and 3T scanners of different models of GE Medical Systems, Philips Medical systems, and SIEMENS at over 33 sites in 11 countries. We used 222 T1-weighted scans from 1.5T scanners and 778 from 3T.
- **HCP**: The Human Connectome Project (HCP) (Van Essen et al., 2012) is an effort to characterize brain connectivity and function and their variability in young healthy adults aged between 25 and 30 years. We used 897 T1-weighted scans from the initial data release.
- **PREVENT-AD**: The PREVENT-AD (Pre-symptomatic Evaluation of Novel or Experimental Treatments for Alzheimer’s Disease, http://www.prevent-alzheimer.ca) program (Tremblay-Mercier et al., 2014) follows healthy individuals age 55 or older with a parental history of AD dementia. We used T1-weighted 1251 scans from the data released in 2017.
- **IPMSA**: To test the generalizability of the model in a completely independent dataset, we used T1-weighted MRI scans of 1200 patients with secondary progressive multiple sclerosis (SPMS), randomly selected from the from International Progressive Multiple Sclerosis Alliance (IPMSA) study (https://www.progressivemsalliance.org/), scanned on 20 different scanner models (including both 3T and 1.5T) across 195 sites (Dadar et al., 2020).

### 2.2 Automatic registration methods

All 9693 image volumes from the evaluation in (Dadar et al., 2018) were used here for training, validation, and testing. Each volume was registered to stereotaxic space defined by MNI-ICBM152-2009c template (referred to below as the MNI template) (Fonov et al., 2011; Manera et al., 2020) using the seven stereotaxic registration procedures listed below yielding a total of 64476 images that completed processing and provided QC images:

- **MRITOTAL (two versions)**: a hierarchical multi-scale 3D registration technique for the purpose of aligning a given MRI volume to an average MRI template aligned with the Talairach stereotaxic coordinate system (Collins et al., 1994). We have tested two configurations of this method: “standard” and “icbm”. The source code is available at https://github.com/BIC-MNI/mni_autoreg.
- **BestLinReg:** a 5-stage hierarchical technique similar to MRITOTAL that is part of the MINC tools and is based on a hierarchical non-linear registration strategy developed by Robbins et al. (Robbins, 2004). We tested two versions: one where cross-correlation coefficient is used as a cost function and another using normalized mutual information. The source code is available at https://github.com/BIC-MNI/EZminc/blob/ITK4/scripts/bestlinreg_s.
- **Revised BestLinReg (two versions)**: This method is the same as BestLinReg above, but with a different set of parameters and normalized mutual information cost function. The source code is available at https://github.com/BIC-MNI/EZminc/blob/ITK4/scripts/bestlinreg_claude.pl. A modified version of Revised BestLinReg using a different set of parameters was also applied.
- **Elastix**: an intensity-based registration tool (Klein et al., 2010). Elastix has a parametric and modular framework, where the user can configure different components of the registration. We used Mattes mutual information, adaptive stochastic gradient descent optimizer with Similarity Transform with 7 parameters https://bitbucket.org/bicnist/bic-nist-registration/src/5d253993e7b9/Elastix.
- **ANTS**: a popular registration tool (Avants et al., 2011) that can function both for linear and nonlinear registration. For this paper we used ANTS for linear registration with following parameters: registration was done in three stages, at first stage only translation was estimated, at the second stage rigid body transformation and finally full affine transformation, each stage used Mattes mutual information as cost function and three hierarchical levels of fitting with different resolution and blurring kernel (4×2×1vox downsampling, and 3×2×1 blurring kernel) using stochastic gradient descent optimizer with gradient step of 0.1.

### 2.3 Manual quality control method

We used the manual QC result from 64476 linear registrations from our previous paper (Dadar et al., 2018) for training, validation and testing. Resulting transformations were stored as affine 4×4 matrices and applied to the corresponding original scans to resample them on a 1mm^3^ voxel grid in stereotaxic space. Then, a series of slices were extracted, and the outline of the MNI template brain was overlaid on top to create an image that was given to the human rater to assess registration performance (See Fig. 1). Assessment was done on a PASS/FAIL basis, blind to the registration method, dataset, and clinical diagnosis.

**Figure 1.**
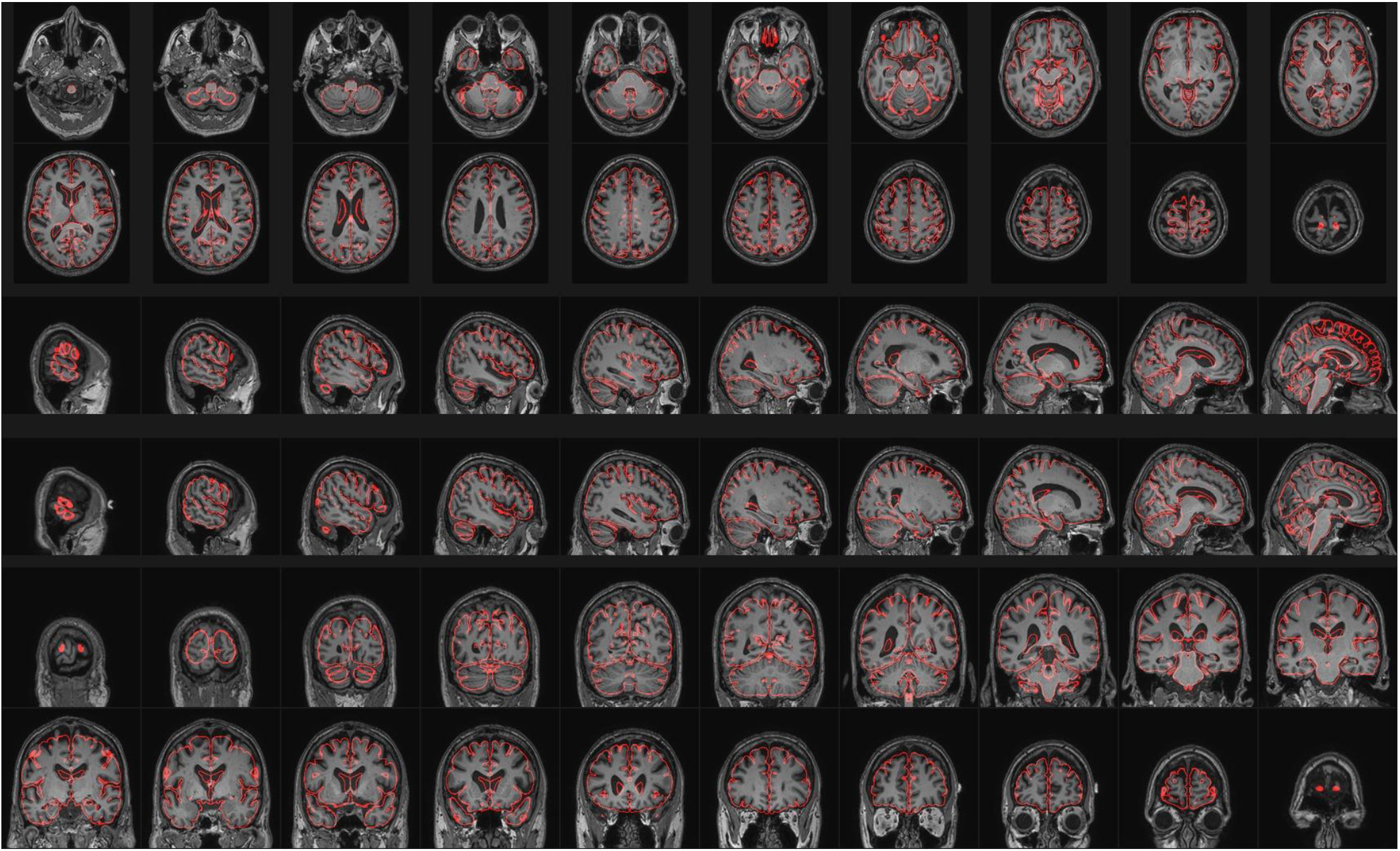
QC image for the human rater. Grayscale - one example subject’s MRI scan after registration and resampling into stereotaxic space on a 1mm isotropic grid, red line - outline of the MNI-ICBM152-2009c brain template. One can see how the average template outline roughly fits the brain contour of the subject indicating a good registration.

Out of 64476 examples, 54458 (**84.5**%) were accepted (passed) and 10018 (**15.5**%) were rejected (failed) by the human rater. To test reproducibility of the human rater results, a random subset consisting of 1000 examples were re-evaluated by the same rater, resulting in intra-rater Dice kappa agreement rate (Sørensen, 1948) of **0.96** and intra-rater agreement rate of **93%**.

### 2.4 Automatic registration quality control with deep neural network

We modified an existing deep neural network (DNN) design to automatically assess stereotaxic registration quality, reusing weights trained on the ImageNet database (Fonov et al., 2018; Gross and Wilber, 2016; Russakovsky et al., 2015). Our motivation was to be able to use a standard pre-trained deep learning model, as a feature extractor, but to make it sensitive to the orientation and position of the object inside the image, as well as combining features from several different slice views.

We compared the performance of several variants of models based on ResNET available in the pytorch library (ResNET-18, 34, 50, 101, 152) (Gross and Wilber, 2016; He et al., 2016). Instead of feeding a single image with 60 different sub-images showing coronal, sagittal and transverse slices of the registered volume as in the human quality control process, we used a simpler set of images that was created in the following fashion: (i) the original 3D MRI scan in the native space, without any preprocessing, was resampled to the MNI template space using the linear transformation matrix provided by registration algorithms; (ii) the whole range of the image intensities of the input file was mapped to the 0.0-1.0 range; (iii) one Axial, one Sagittal and one Coronal slice were extracted from the middle of the registered 3D volume; (iv) the three 2D slice images were resampled to have 256 pixels in the longest dimension and then cropped around the central area to 224×224 pixels. These images were stacked to form a 224×224×3 dataset that was used as input to the model. Figure 2A shows an example of three images corresponding to a scan that passed manual QC and three others that failed QC. Figure 2C shows the input of DNN when reference image of the MNI template is used.

**Figure 2.**
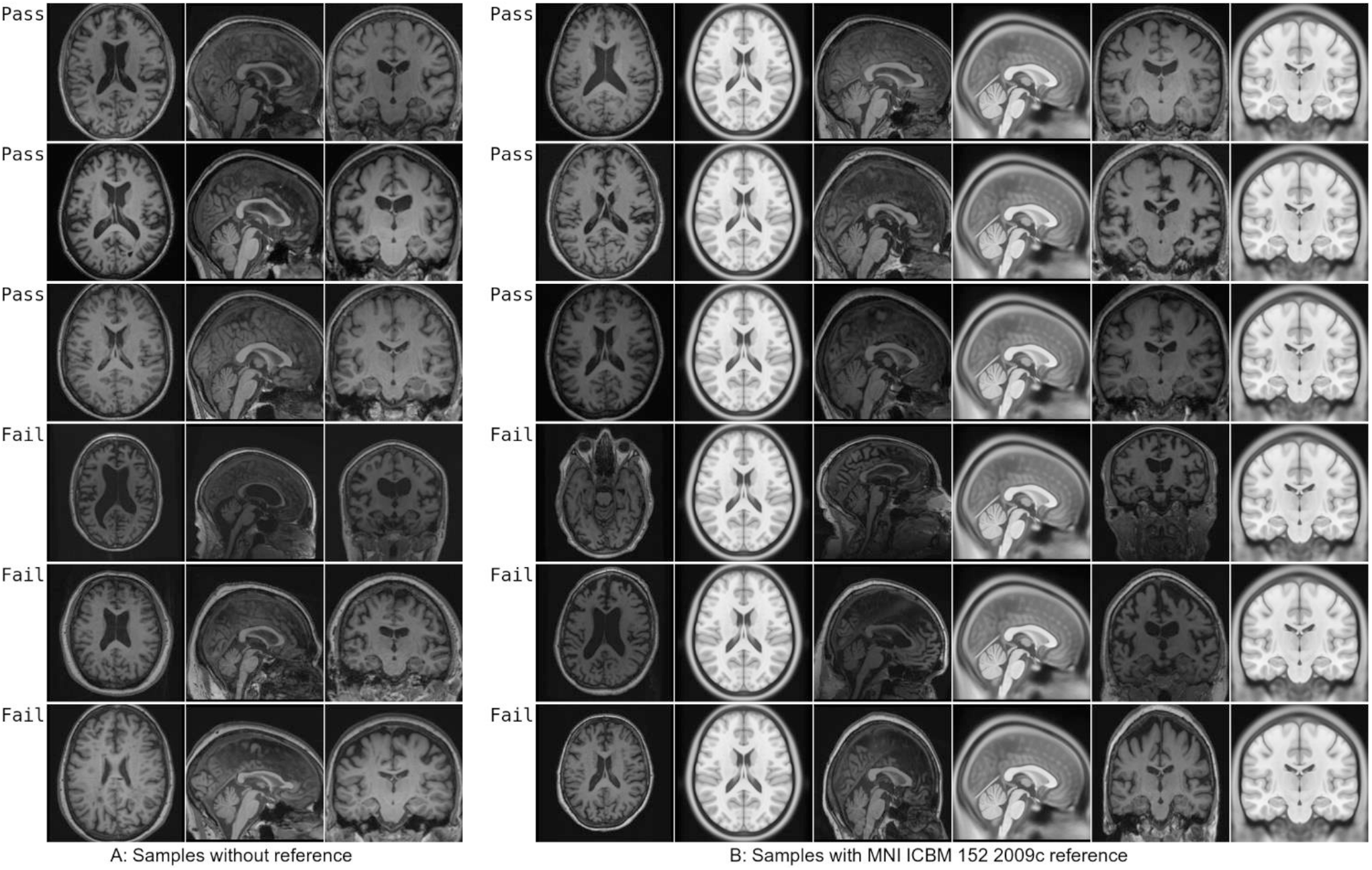
Examples of images generated for automated QC script, used as input for QCResNET and DistResNET training

In order to transfer domain knowledge, we modified a DNN trained on the ImageNet dataset (DNN0) in the following fashion (see Fig. 3): (i) the input layer was altered to deal with grayscale images by collapsing the weight tensor along the dimension corresponding to input RGB dimension; (ii) the last few layers corresponding to high level features used for ImageNet classification and spatial pooling were removed; (iii) each input feature (i.e channel) from the image stack was processed sequentially using the same DNN0 layer; (iv) outputs of DNN0 layer corresponding to the three different images from the stack were concatenated and used as inputs to the last several layers, replacing the behavior of the layers originally removed from DNN0 with the goal of making them more sensitive to spatial orientation and position; (v) the final layer was modified to produce one of two types of output; a binary pass or fail or a continuous scalar estimate of misregistration distance. The resulting model is called QCResNET-X for pass/fail model and DistResNET-X for misregistration distance model, where X corresponds to the choice of underlying ResNET (i.e., ResNET-18, 34, 50, 101, 152).

**Figure 3.**
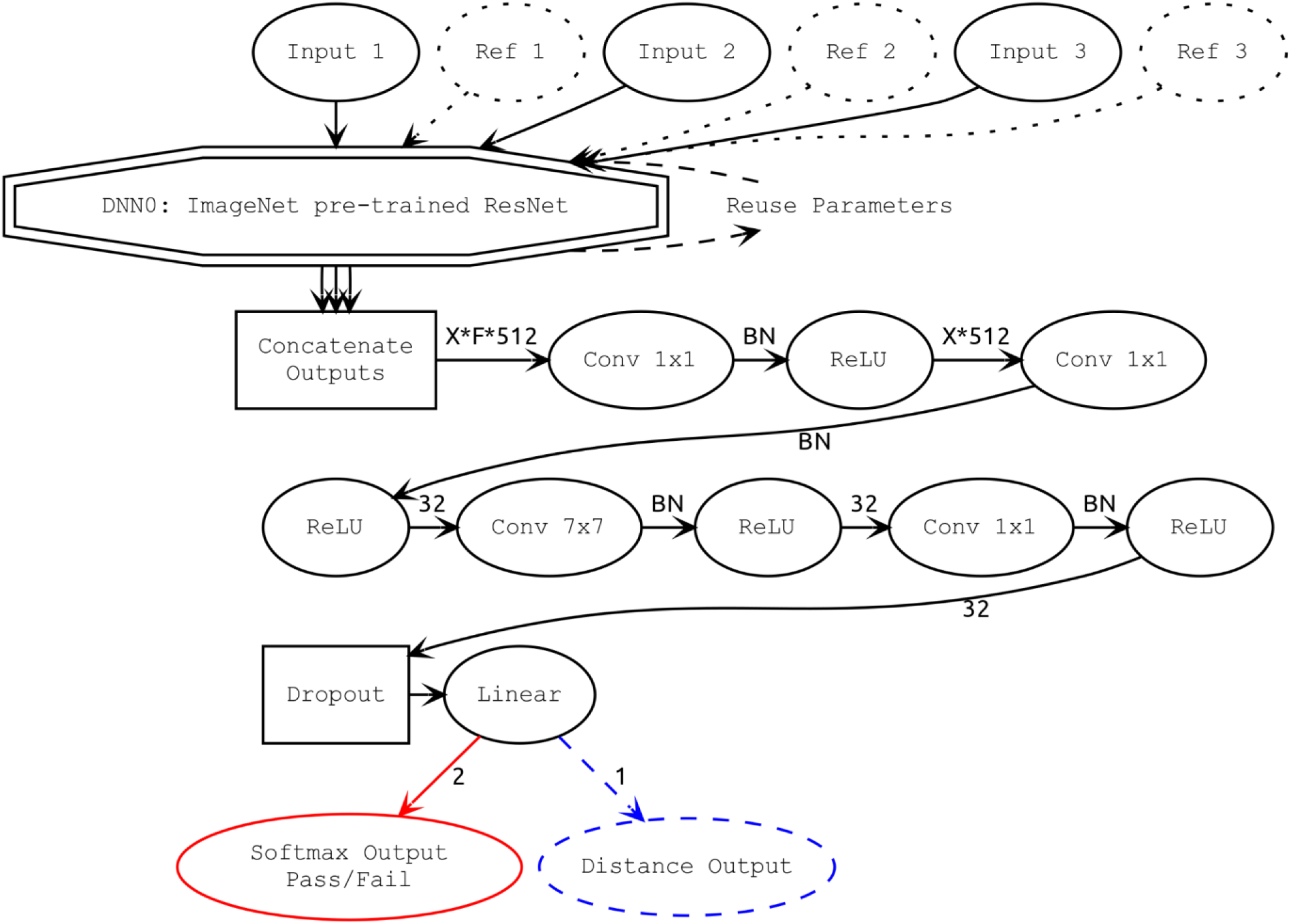
Overall QCResNET-X and DistResNET-X design, dotted nodes represent optional components; BN - batch normalization; ReLU - rectified linear unit; Conv - 2 dimensional convolution; numbers are number of channels, X-ResNET expansion factor, F - number of input features (3 or 6); Red and Blue colored parts are QCResNET and DistResNET evaluation outputs respectively.

We also created an alternative scheme, where reference images extracted from MNI template (Fig. 2B) were used as an additional set of features in the image stack. In this case, the model was modified to accept 6 input grayscale feature maps – combining those that were subject-specific, and the others that were fixed for all subjects (see Fig. 2B) In this scheme, the network input was six images stacked together, to form 224×224×6 dataset in the following order Axial sample, Axial Reference, Coronal Sample, Coronal Reference, Sagittal sample, Sagittal Reference.

We compared two approaches for misregistration representation: one based on binary classification (i.e pass or fail) and the second one similar to the method from (de Senneville et al., 2020) where misregistration distance is estimated. Cross-entropy loss function was used as an objective function in pass/fail classification and mean square error objective function in case of distance estimation. The ADAM optimizer was used to train the network.

### 2.5 Silver standard and distance estimation

To estimate misregistration distance, we created a “silver standard” transformation for each original MRI volume, by averaging all transformations that passed manual QC (i.e., averaging up to six transforms for each subject using the six methods described in section 2.2). We then calculated the distance between each estimated transformation and the “silver standard”, as defined in eq (1), where ROI_icbm_ is the bounding box of the brain ROI of the ICBM 152 2009c template (Fonov et al., 2011; Manera et al., 2020), X_a_ - silver standard transform, X_b_ - estimated transform, distance is expressed in millimeters in stereotaxic space.

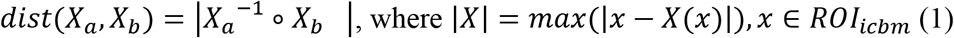

### 2.6 Data augmentation

#### 2.6.1 Pass/Fail classification model

For the purpose of training the model for pass/fail approach we created additional QC images, based on the information from manually QCed results. To simulate image acquisition with thick slices, the preregistered scans were downsampled in z direction using scaling factor randomly chosen between 1, 2 and 3; to simulate a restricted field of view, we randomly cropped either top 20% slices, or bottom 20% of slices or none; to simulate small imperfections of the registration parameters we added random rotations around x, y, and z axis in the range [-0.1,0.1] degrees; and finally, a random flip along the x (left-right) axis was added. We generated 5 augmented samples per each original sample, resulting in 322460 samples. We used the same manual QC labels for the augmented datasets, with the assumption that the data augmentation steps applied to the preregistered data change the characteristics of the data, and because we change the registration parameters only slightly, these augmentation steps do not change the outcome of the manual stereotaxic registration QC.

#### 2.6.2 Distance estimation model

In case of the distance estimation training, we used an approach similar to the one published in (de Senneville et al., 2020): we used only samples where manual quality control was passed, and then generated random transformations with uniform distribution of distances from the “silver standard” (see below). We also simulated random flips along x, and random cropping in z direction to simulate restricted field of view and downsampling factor in z direction. We produced 65275 training samples in this approach.

### 2.7 Network training

We used an 8-fold cross-validation scheme, where all available samples are split into 8 equal partitions, based on unique subjects. At each round of cross-validation, the corresponding registration result of each partition is used as “testing” dataset and the rest of data is used as “training” dataset. A small subset of the “training” dataset (200 unique subjects) was excluded from training and used for on-line validation (“validation” dataset). To compute statistics, the “validation” and “testing” datasets were balanced to have an equal mix of pass/fail samples. Training was performed in epochs when all available samples from the “training” subset were passed through the training loop. At the end of each epoch, the performance of the current model was evaluated using the “validation” dataset and the four models achieving best performance in terms of true negative rate, true positive rate, accuracy (ACC) and area under the ROC curve (AUC) were preserved.

An initial experiment to estimate the number of epochs needed to train a classification model, we compared the performance of QCResNET 18,34,50,101,152 after 30 epochs, using only one fold out of 8. Performance on the validation dataset was evaluated after every 200 mini-batches; accuracy, true positive rate, true negative rate and area under the ROC curve were calculated. Final performance of each model was assessed on the “testing” dataset, and only AUC was calculated. Based on the results shown in Figure 4, we determined that training 10 epochs was enough for all models. Note that, for example, the model achieving best TNR for QCResNET18 was found after 10^th^ epoch, and the best accuracy after 4^th^ epoch.

**Figure 4.**
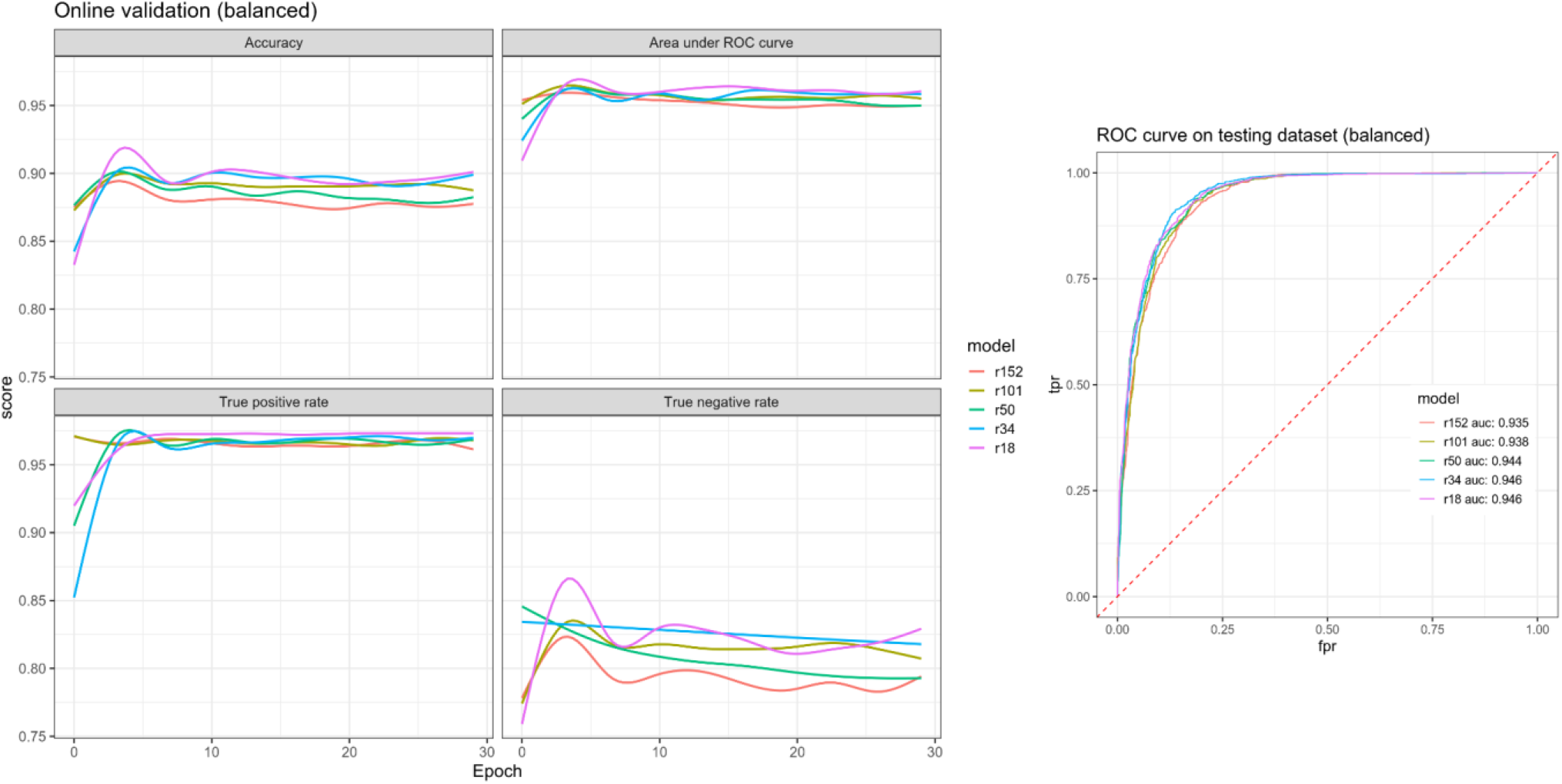
Progressions of training for QCResNET-18,34,50,101,152 with reference. Left: performance on the online validation dataset calculated after each 200 mini-batches, smoothed with loess function; Right:AUC of the performance on testing dataset. Overfitting is visible after approximately 10 epochs. QCResNET 18 and 34 shows the best performance in terms of AUC.

As a result of the initial experiment performed to determine the optimal number of epochs used for training, we noticed that the final performance depends on which epoch was chosen for early stopping to avoid overfitting. We therefore also compared the performance of the method based on resnet-18 using 8-fold cross validation, depending which metric was used for early stopping (Figure 5). The model achieving the highest TNR, is also guaranteed to achieve the lowest FPR, since FPR=1-TNR. This model was chosen as a final model for the training.

**Figure 5.**
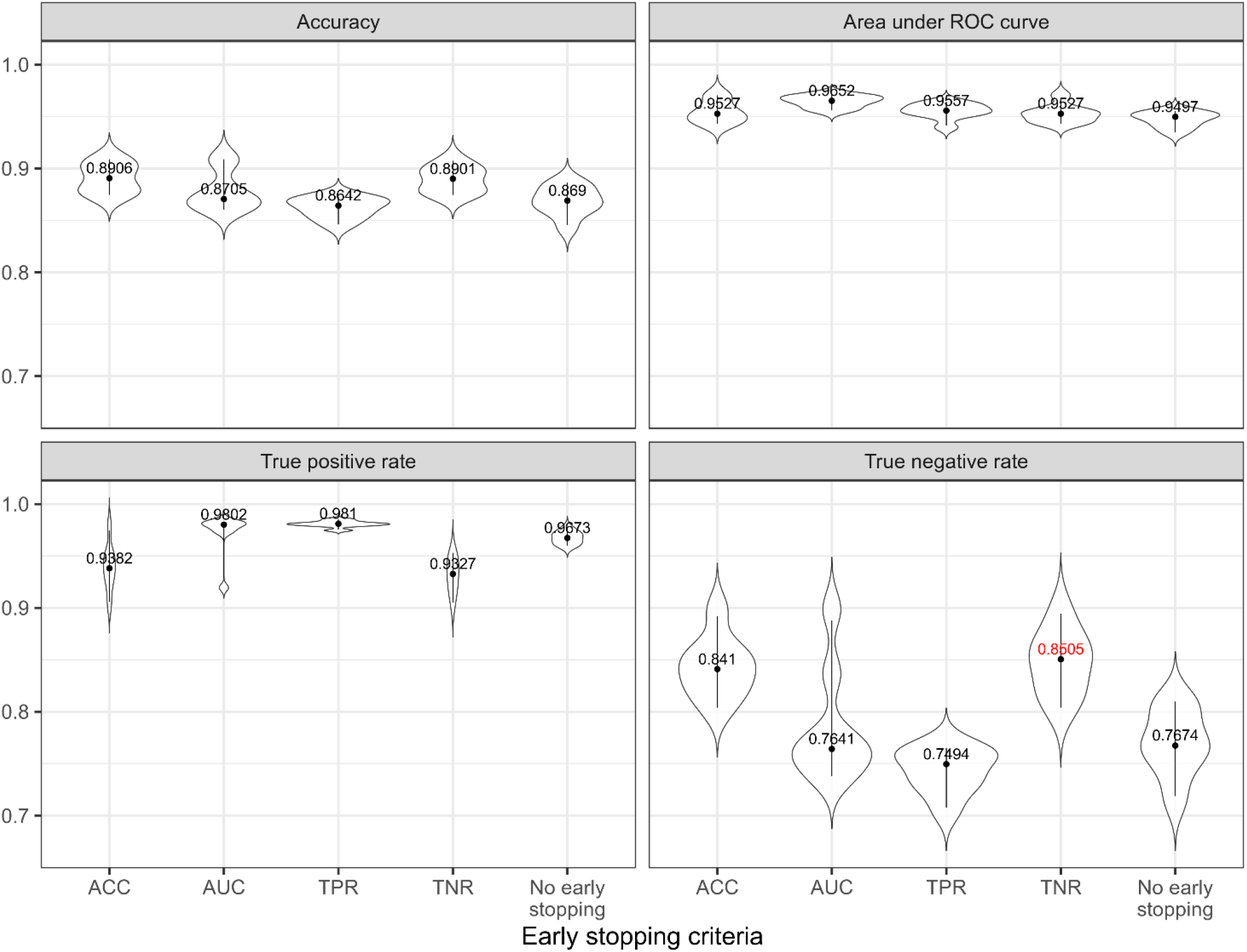
Violin plots showing performance of the QCResNET-18 with reference, depending on the early stopping criteria. Vertical line represents interquartile range, black dot – median value. Value highlighted in red corresponds to the model achieving the best true negative rate.

Other training considerations: we used gradient magnitude clipping at 1.0 to avoid the problem of exploding gradients, learning rate of 1e-5 was used for all models, 100 warm up iterations with learning rate of 1e-9 were used to initialize weights of ADAM optimizer.

In the case of the distance estimation model, we trained a model DistResNET-18 for 30 epochs, calculating mean squared error in distance estimation after each 200 mini-batches. We used only augmented samples for training the model, and samples from the original manual QC database for testing to mimic, as closely as possible, the training regimen of (de Senneville et al., 2020). Figure 6 shows the progression and the result of training for 1 fold out 8.

**Figure 6.**
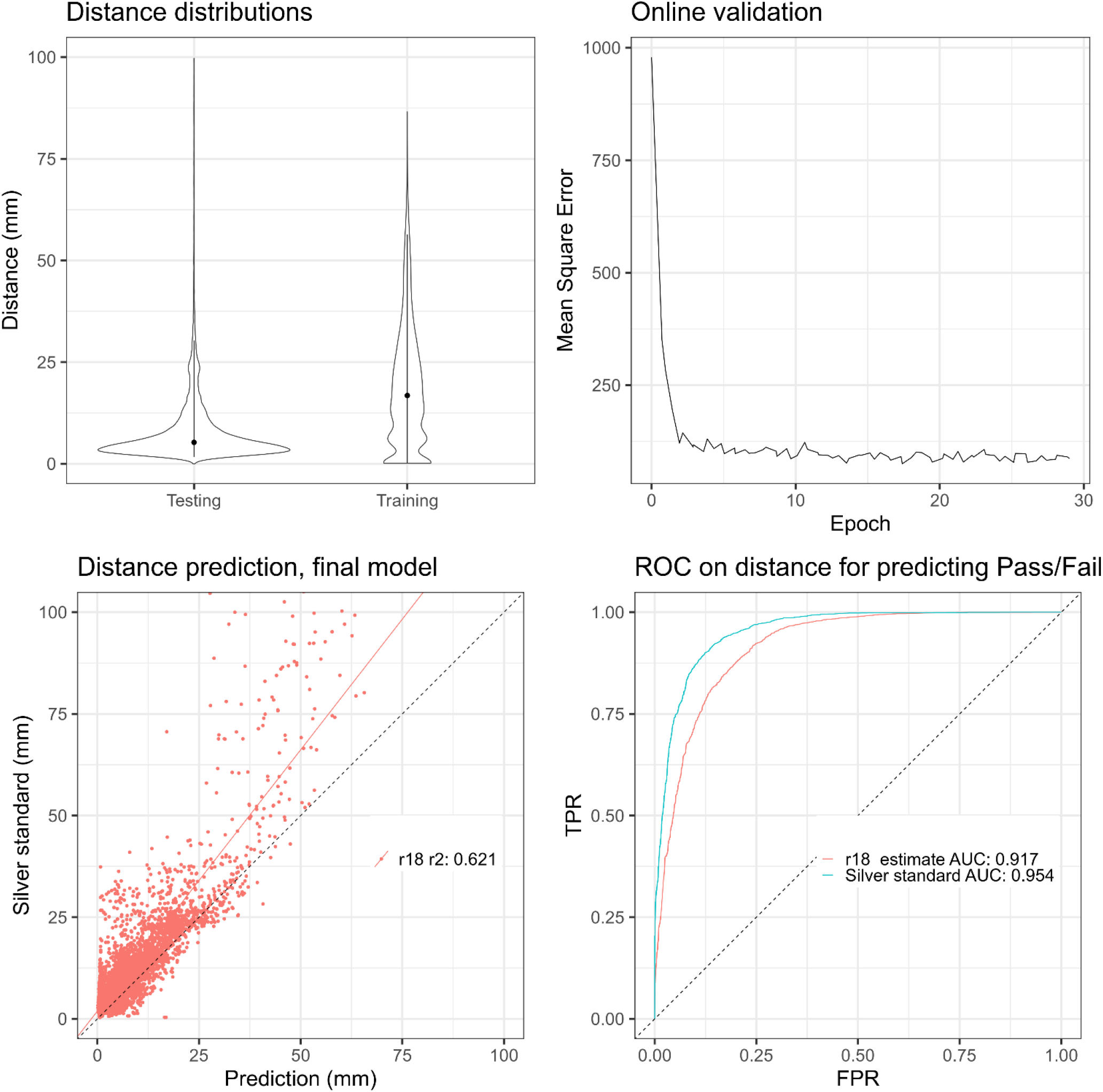
Training distances estimation model (DistResNET-18), 30 epochs, mean squared error is calculated after 200 mini-batches.

To test the final performance of DistResNET-18, 34, 50, 101, 152, we compared the predicted misregistration against the silver standard, in terms of root mean square difference. We also calculated the area under the curve for each experiment. We used early stopping based on the means squared distance from the ground truth for the validation dataset.

### 2.7. Independent sample validation

To assess the performance of the proposed method on an independent sample not used during the training and cross-validation experiments, we applied the revised_bestlinreg (Dadar et al., 2018) linear registration technique to 22,757 scans from the IPMSA dataset. Manual QC was performed sequentially until 600 samples that passed QC and another 600 samples that failed QC were identified. Then these samples were used to test performance of the DARQ QCResNet-18 and DistResNet-18.

### 2.8 Data and Code Availability Statement

The data that support the findings of this study were obtained from the ADNI (publicly available at http://adni.loni.usc.edu/), PPMI (publicly available at https://www.ppmi-info.org/), HCP (publicly available at https://www.humanconnectome.org/), and PREVENT-AD (publicly available at https://portal.conp.ca/dataset?id=projects/preventad-open) datasets. IPMSA is not publicly available. The original implementation (Fonov et al., 2018) was done using the Torch library (Collobert et al., 2011), and we have re-implemented the software in pyTorch (Ketkar, 2017). All experiments were performed using the Torch version. The source code of the method, implemented in torch and pyTorch (python) and pre-trained neural network is publicly available at https://github.com/vfonov/DARQ

## 3 Results

Following the initial experiment with 30 epochs mini-batches using one out of 8 folds, we observed over-fitting after 10 epochs, so the rest of cross-validation experiments were conducted with 10 epochs.

We tested DARQ models QCResNET-18, −34, −50, −101, and −152. Two training schemes were tested: 1. using only QC images (No Reference) and 2. combining QC images with the images of the reference dataset, i.e. MNI template (With Reference). The resulting agreement with the manual rater measured by accuracy, true positive rate, true negative rate and area under receiver operating characteristic curve (Robin et al., 2011) were used to evaluate performance of the models; we used early stopping criterion based on the best TPR. The QCResNET-18 with reference images showed the best results in terms of TNR, at the expense of slightly lower TPR (see Figure 7).

**Figure 7.**
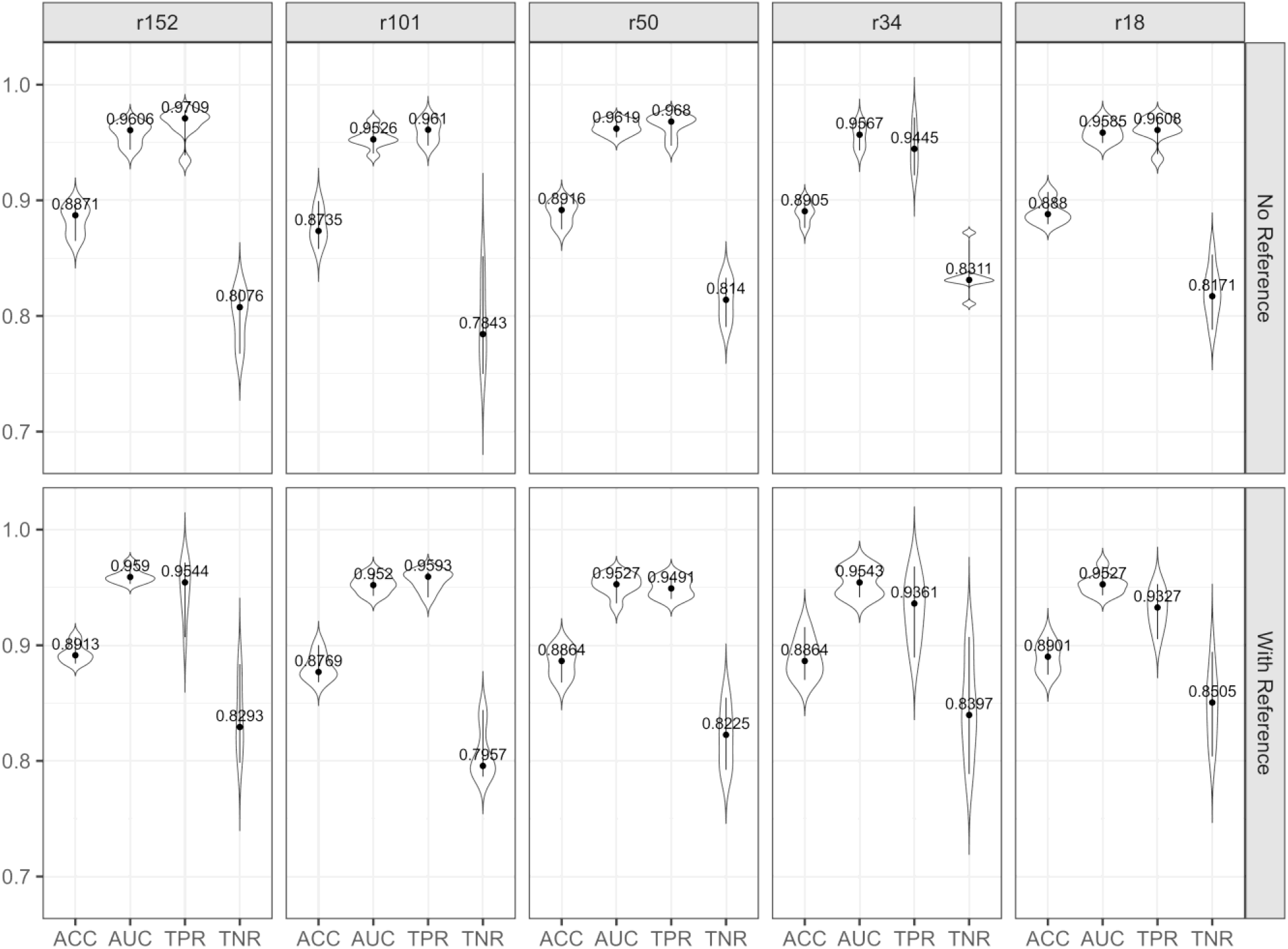
Violin plots showing performance of all tested classification models with and without reference image, compared to the manual rater.

Results for experiment with Distance Evaluation model (DistResNet-18, 34, 50, 101, 152) are shown in Figure 8.

**Figure 8.**
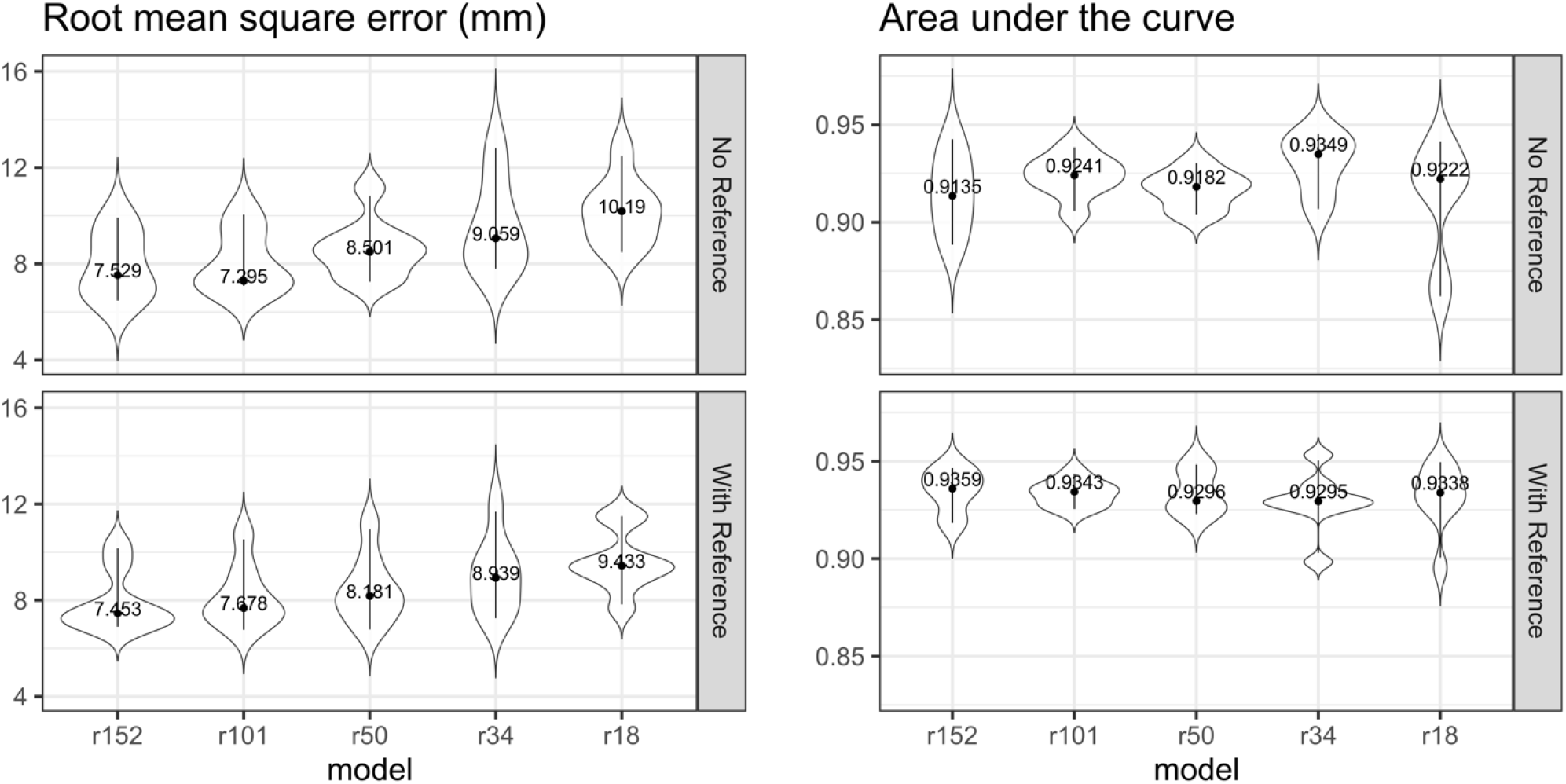
Performance of all tested distance estimation models with and without reference image

As expected, the more complex model (i.e., DistResNet-152) estimated the misregistration distance better. On the other hand, if the distance were to be used for classification task, based on a threshold, the auc analysis shows that all models with reference showed approximately the same result.

Comparing performance of the DistResNet vs QCResNet, all distance estimation models achieved lower AUC than classification models (see Fig. 7,8), in particular the best AUC for DistResNet152 with reference is 0.9359 and the AUC for QCResNet152 with reference is 0.9590 for the model with equal number of parameter (statistical significance of the difference p=0.0002 using Wilcoxon rank sum exact test)

Finally, we only used the QCResNET-18 model with reference (i.e., the model with the best performance based on the previous assessments) for the distance experiments. The distribution of the distances between “silver standard” transformations and transformations estimated by each method is shown on Figure 9 (left panel). The behaviour of the distances depending on the automated QC outcome is also shown on Figure 9 (right panel). For the manual quality control, the cases that passed had distances (mean±sd) values of 5.96±2.93 mm and those that failed had values of 25.2±23.2 mm. For the automated QC, true positive cases had the lowest average distance compared with the silver standard 5.68±2.32 mm, followed by false negative 9.7±5.36 mm, false positive 12.2±6.6 mm, and true negative 27.6±24.4 mm cases.

**Figure 9.**
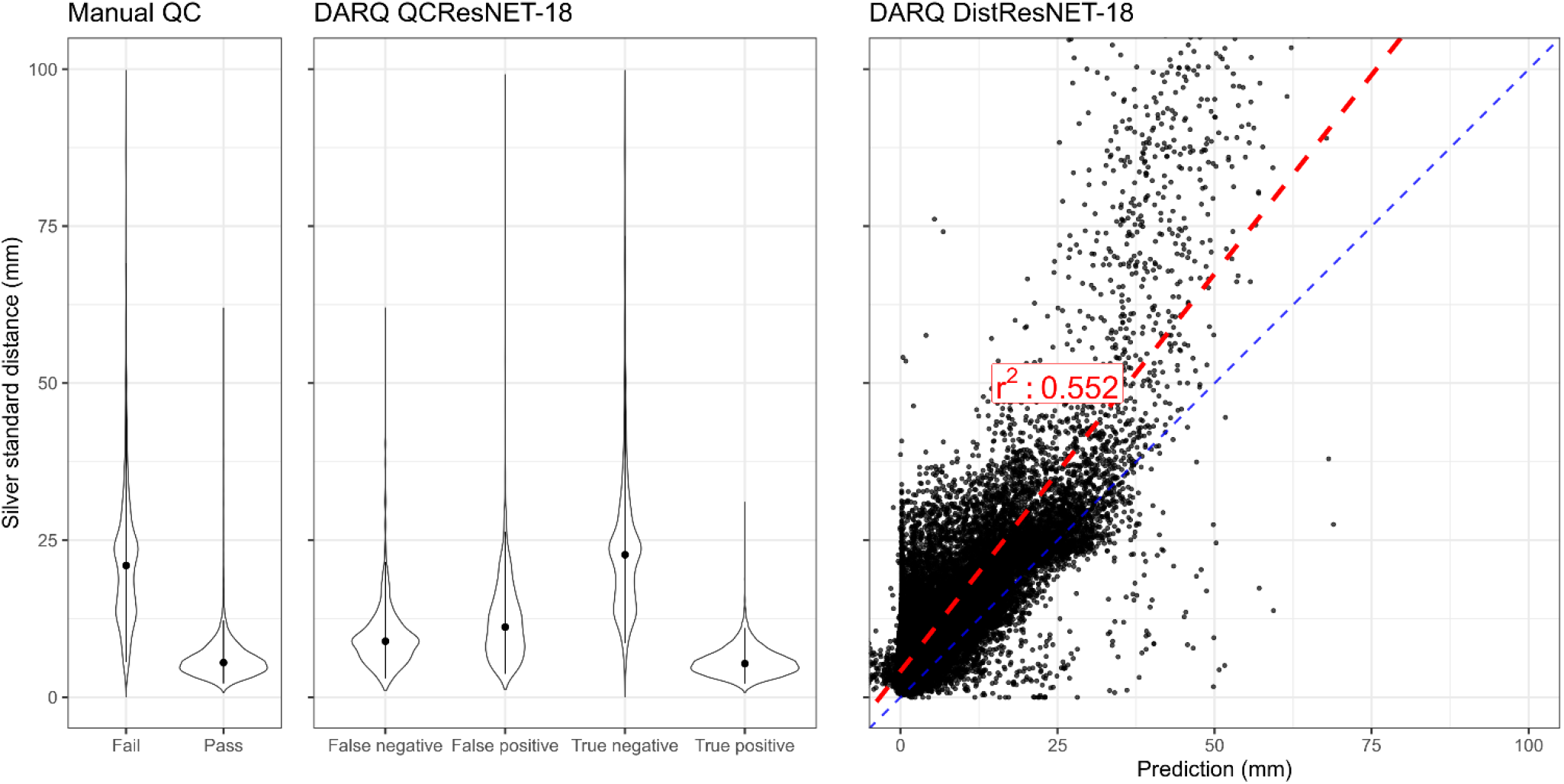
Distance from the “silver standard” in mm, for the manual QC results, and for the outcome of DARQ QCResNet-18 and DistResNet-18 with reference methods cross-validation.

To assess the generalizability of DARQ to independent data with different acquisition parameters and from a variety of scanner models and manufacturers, we applied QCResNET-18 to the multi-center and multi-scanner IPMSA dataset and achieved ACC = 96.1%, TPR= 96.7%, TNR = 95.5%, AUC = 98.7%. Figure 10 shows the distribution of the continuous output generated by the automated QC tool for the passed and failed registrations for the IPMSA data.

**Figure 10.**
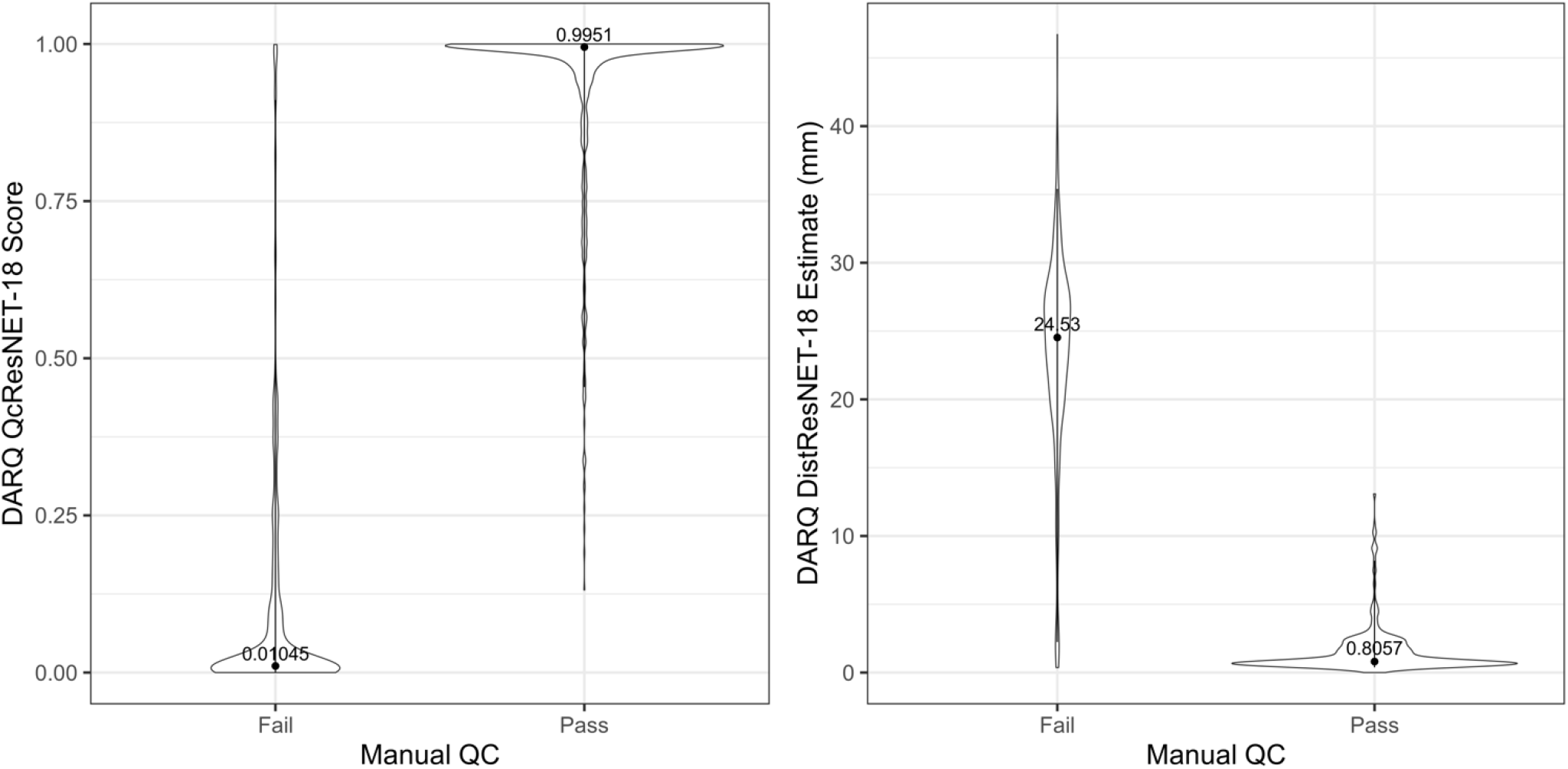
Distribution of the continuous outputs generated by the automated QC tool for the registrations passed and failed by the human rater.

## 4 Discussion

In this paper, we proposed a deep learning based, fully automated tool, for quality control of linear stereotaxic registration. We have demonstrated that it is possible to automate the task of manual quality control using a deep learning network with results comparable to the human rater. The proposed tool had good agreement with the human ratings in the cross-validation experiments as well as when tested on an independent sample not used for training.

Note that not all stereotaxic registration that FAIL QC are clear catastrophic failures. Manual QC was evaluated using QC images like that shown in Figure 1 and the rater could zoom into different parts of the composite image to make their decision. Since linear registration only accounts for the global parameters of translation, rotation and scaling, there will be different levels of local residual misregistration between the grey level image of the subject and the red contours of the template. During manual QC, these differences drive the subjective decision made to PASS or FAIL a dataset. For example, some cases have a z-scale that is slightly too large in the opinion of one rater, while for another rater, the curvature at the apex of the brain might be considered to be simply due to residual misalignment between subject and template and thus rated as a PASS, thus giving rise to inter-rater agreement that is not quite equal to 1.0.

All automatic models had excellent performances in detecting failed registration cases, with accuracies ranging between 87.3% to 89.1% with balanced testing sets. QCResNET-18 network with reference had the best performance in terms of TNR, and was selected as the final model. Interestingly, all methods had better performances in terms of false positive rate (Fig. 7) when reference images extracted from the MNI template were also used as an additional set of features.

The performances of the deep learning methods (with accuracy values ranging between 87.3-89.1% for the cross-validation experiment and 96.1% for the independent sample) were comparable with the intra-rater variability (test-retest accuracy of 93%), indicating that some of the disagreements might be due to human rater variability.

To enable a more quantitative assessment of the registrations, we defined a silver standard transformation for each scan, based on the average of all transformations that passed QC by the human rater. As expected, the distance between this silver standard and passed registrations was much lower 6.6 (4.5) mm than the failed registrations for the manual ratings 28.1±31.4 mm. Similarly, the distance for the true positive automated QC results was also much lower 6.3±4.1 mm than the true negative automated QC results 32.0±34.3 mm. Interestingly, the average distance for both false negative 11.5±8.8 mm and false positive 15.3±12.0 mm cases was somewhere in between (Fig. 8), indicating that those are likely borderline cases that might have been failed by a stricter human rater. This was also our impression when we visually assessed the cases passed in the manual QC and failed by DARQ and vice versa. Evaluation of the performance of the proposed method on an independent multi-center and multi-scanner dataset showed that the results are generalizable to data with different acquisition protocols obtained from different scanner models and manufacturers. Compared with the manual rater, DARQ had excellent FPR (1.8%), correctly identifying the failed registrations.

We also compared our classification approach to the misregistration estimation approach, similar to that used by de Senneville et al. (de Senneville et al., 2020). Based on our experimental results, while the DistResNet misregistration model was able to estimate distances (compared against our silver standard distance), it achieved a lower performance than our proposed technique in terms of (pass/fail) registration classification (auc values ranging from 91.3 to 93.5 versus 95.2 to 96.2, Figs 7 and 8). For example, a small rotation error, combined with a small scaling error may pass QC, but the same distance with solely a translation error may not be acceptable for QC (Fig. 9). Therefore, while the registration error distance estimation method might be beneficial in other applications, for the specific task of registration QC, our proposed method may be superior.

Since the majority of scans are usually accurately registered by most commonly used registration methods, higher sensitivity rates (i.e., detection of failed scans, as opposed to higher specificity) are particularly desirable for an automated QC tool, since it is much more practical (i.e. less time consuming) for a human rater to verify quality of a small number of false positives (registration that are falsely identified failures) as opposed to a large number of false negatives (good registrations that are identified failed scans). For example, out of 22,757 registrations from the IPMSA dataset, the automated QC tool failed only 930 cases (4.09%). If we are confident that all the failed cases are accurately captured by the automated QC tool, the human rater only needs to assess the quality of the very small percentage of cases that were failed by the automated QC tool. In addition, removing human effort from (most of) the registration QC process enables the users to rerun the registration methods for the failed cases with different settings and parameters to ensure a high acceptance rate for their cohort of interest. In other words, the proposed QC procedure could be incorporated into the process of the registration to enforce the method to repeat registration with different parameter settings until an accepted registration (by the automated QC tool) is obtained. A false negative (failing an acceptable registration) in this case would only increase registration time by forcing the method to repeat the process. Therefore, a lower TPR at the expense of a low FPR would be tolerable and lead to the overall improvement of the registration performance. The use of such automated QC tools may likely affect the outcome of processing pipelines. This is a complex and open issue as the answer is dependent on the specific pipeline and research question and should be the subject of future work. Other automated QC registration tools are built on simulated data (de Senneville et al., 2020). Instead of artificially generating failed registrations (i.e., by applying randomly generated incorrect transformations to the original images), we trained and validated the performance of DARQ on a four large multi-center and multi-scanner datasets comprised of actual registrations that were produced by different commonly used image registration pipelines. This enables the proposed method to learn the realistic types of registration failures that commonly occur, as opposed to synthetically generated failures that might be much easier to capture, but might not be likely to occur in real settings.

We have shown that DARQ is robust over a wide range of ages and different disease states (i.e., healthy, AD, PD, MS). We note that the ICBM152_2009c template used here is defined to be in the same space as the other non-linear unbiased average ICBM152 templates (found at http://nist.mni.mcgill.ca/icbm-152-nonlinear-atlases-2009) and the linear average ICBM152 template (found at http://nist.mni.mcgill.ca/icbm-152lin) as well as the older MNI305 template (http://nist.mni.mcgill.ca/mni-average-brain-305-mri). A stereotaxic transformation to the ICBM152_2009c template that passes QC would also pass QC for registration to these other templates as there are almost no differences in position, orientation or extent in the left-right, AP or SI. In future work, it may be interesting to apply a strategy similar to that proposed here to develop new tools to automatically QC other registration tasks, e.g., linear registration to other target images, multi-modality registrations, and non-linear stereotaxic registrations. The open-source implementation of DARQ that we have made publicly available will allow researchers to adapt and retrain our proposed model to perform a wider range of QC tasks once a suitably large set of training data becomes available.

Manual quality control of linear registrations is time consuming (~30 hours for 9693 registrations) and prone to inter-rater and intra-rater errors (Dadar et al., 2018), making it particularly challenging for large databases such as the UK Biobank, NACC, and ADNI. DARQ is generalizable to data from different scanners since it has been trained and extensively validated on data from multiple scanners. The proposed technique will save a significant amount of human effort in processing large imaging databases and will increase reproducibility of results. Finally, the proposed automated QC tool could be used for fast and efficient automatic testing of the robustness of image registration methods. Finally, we have made two implementations of DARQ along with the pre-trained models publicly available.

## 5 Acknowledgements

We would like to acknowledge funding from the Famille Louise & André Charron. MD is supported by a scholarship from the Canadian Consortium on Neurodegeneration in Aging in which DLC is a co-investigator as well as an Alzheimer Society Research Program (ASRP) postdoctoral award. The Consortium is supported by a grant from the Canadian Institutes of Health Research with funding from several partners including the Alzheimer Society of Canada, Sanofi, and Women’s Brain Health Initiative.

Data collection and sharing for this project was in part funded by the Alzheimer’s Disease Neuroimaging Initiative (ADNI) (National Institutes of Health Grant U01 AG024904) and DOD ADNI (Department of Defense award number W81XWH-12-2-0012). ADNI is funded by the National Institute on Aging, the National Institute of Biomedical Imaging and Bioengineering, and through generous contributions from the following: AbbVie, Alzheimer’s Association; Alzheimer’s Drug Discovery Foundation; Araclon Biotech; BioClinica, Inc.; Biogen; Bristol-Myers Squibb Company; CereSpir, Inc.; Cogstate; Eisai Inc.; Elan Pharmaceuticals, Inc.; Eli Lilly and Company; EuroImmun; F. Hoffmann-La Roche Ltd and its affiliated company Genentech, Inc.; Fujirebio; GE Healthcare; IXICO Ltd.; Janssen Alzheimer Immunotherapy Research & Development, LLC.; Johnson & Johnson Pharmaceutical Research & Development LLC.; Lumosity; Lundbeck; Merck & Co., Inc.; Meso Scale Diagnostics, LLC.; NeuroRx Research; Neurotrack Technologies; Novartis Pharmaceuticals Corporation; Pfizer Inc.; Piramal Imaging; Servier; Takeda Pharmaceutical Company; and Transition Therapeutics. The Canadian Institutes of Health Research is providing funds to support ADNI clinical sites in Canada. Private sector contributions are facilitated by the Foundation for the National Institutes of Health (www.fnih.org). The grantee organization is the Northern California Institute for Research and Education, and the study is coordinated by the Alzheimer’s Therapeutic Research Institute at the University of Southern California. ADNI data are disseminated by the Laboratory for Neuro Imaging at the University of Southern California.

Data used in this article were in part obtained from the Parkinson’s Progression Markers Initiative (PPMI) database (www.ppmi-info.org/data). For up-to-date information on the study, visit www.ppmi-info.org. PPMI is sponsored and partially funded by the Michael J Fox Foundation for Parkinson’s Research and funding partners, including AbbVie, Avid Radiopharmaceuticals, Biogen, Bristol-Myers Squibb, Covance, GE Healthcare, Genentech, GlaxoSmithKline (GSK), Eli Lilly and Company, Lundbeck, Merck, Meso Scale Discovery (MSD), Pfizer, Piramal Imaging, Roche, Servier, and UCB (www.ppmi-info.org/fundingpartners).

HCP data was obtained from the Human Connectome Project, WU-Minn Consortium (Principal Investigators: David Van Essen and Kamil Ugurbil; 1U54MH091657) funded by the 16 NIH Institutes and Centers that support the NIH Blueprint for Neuroscience Research; and by the McDonnell Center for Systems Neuroscience at Washington University.

PREVENT-AD data were obtained from the Pre-symptomatic Evaluation of Novel or Experimental Treatments for Alzheimer’s Disease (PREVENT-AD, http://www.prevent-alzheimer.ca) program data release 3.0 (2016-11-30). Data collection and sharing for this project were supported by its sponsors, McGill University, the Fonds de Research du Québec – Santé, the Douglas Hospital Research Centre and Foundation, the Government of Canada, the Canadian Foundation for Innovation, the Levesque Foundation, and an unrestricted gift from Pfizer Canada. Private sector contributions are facilitated by the Development Office of the McGill University Faculty of Medicine and by the Douglas Hospital Research Centre Foundation (http://www.douglas.qc.ca/).

## Notes

https://github.com/vfonov/DARQ

### Competing Interest Statement

The authors have declared no competing interest.

### Summary of Updates

Paper was updated to answer questions raised by peer review: 1. Figures were updated to improve legibility, with box plots replaced by violin plots 2. Acronyms for various measures of similarity were cleaned up and defined only once (i.e true positive rate -> TPR) 3. Description of the figures was improved 4. Larges change was done to the discussion section to clarify areas of applicability of the method and discuss future work.

## References

Alfaro-Almagro, F., Jenkinson, M., Bangerter, N.K., Andersson, J.L., Griffanti, L., Douaud, G., Sotiropoulos, S.N., Jbabdi, S., Hernandez-Fernandez, M., Vallee, E., 2018. Image processing and Quality Control for the first 10,000 brain imaging datasets from UK Biobank. Neuroimage 166, 400–424.

Ashburner, J., Friston, K.J., 2000. Voxel-based morphometry—the methods. Neuroimage 11, 805–821.

Avants, B.B., Tustison, N.J., Song, G., Cook, P.A., Klein, A., Gee, J.C., 2011. A reproducible evaluation of ANTs similarity metric performance in brain image registration. NeuroImage 54, 2033–2044. https://doi.org/10.1016/j.neuroimage.2010.09.025

Benhajali, Y., Badhwar, A., Spiers, H., Urchs, S., Armoza, J., Ong, T., Pérusse, D., Bellec, P., 2020. A Standardized Protocol for Efficient and Reliable Quality Control of Brain Registration in Functional MRI Studies. Front. Neuroinformatics 14, 7. https://doi.org/10.3389/fninf.2020.00007

Canziani, A., Paszke, A., Culurciello, E., 2016. An analysis of deep neural network models for practical applications. ArXiv Prepr. ArXiv160507678.

Collins, D.L., Neelin, P., Peters, T.M., Evans, A.C., 1994. Automatic 3D intersubject registration of MR volumetric data in standardized Talairach space. J. Comput. Assist. Tomogr. 18, 192–205.

Collobert, R., Kavukcuoglu, K., Farabet, C., 2011. Torch7: A Matlab-like Environment for Machine Learning neural information processing systems.

Dadar, M., Fonov, V.S., Collins, D.L., Initiative, A.D.N., 2018. A comparison of publicly available linear MRI stereotaxic registration techniques. NeuroImage 174, 191–200.

Dadar, M., Narayanan, S., Arnod, D.L., Collins, D.L., Maranzano, J., 2020. Conversion of Diffusely Abnormal White Matter to Focal Lesions is Linked to Progression in Secondary Progressive Multiple Sclerosis. Mult. Scler. J. 832345.

de Senneville, B.D., Manjón, J.V., Coupé, P., 2020. RegQCNET: Deep quality control for image-to-template brain MRI affine registration. Phys. Med. Biol. 65, 225022.

Dubost, F., de Bruijne, M., Nardin, M., Dalca, A.V., Donahue, K.L., Giese, A.-K., Etherton, M.R., Wu, O., de Groot, M., Niessen, W., 2020. Multi-atlas image registration of clinical data with automated quality assessment using ventricle segmentation. Med. Image Anal. 63, 101698.

Fischl, B., 2012. FreeSurfer. Neuroimage 62, 774–781.

Fonov, V., Evans, A.C., Botteron, K., Almli, C.R., McKinstry, R.C., Collins, D.L., 2011. Unbiased average age-appropriate atlases for pediatric studies. NeuroImage 54, 313–327. https://doi.org/10.1016/j.neuroimage.2010.07.033

Fonov, V.S., Dadar, M., Collins, D.L., Group, P.-A.R., 2018. Deep learning of quality control for stereotaxic registration of human brain MRI. bioRxiv 303487.

Gross, S., Wilber, M., 2016. Training and investigating residual nets. Facebook AI Res. 6, 3.

He, K., Zhang, X., Ren, S., Sun, J., 2016. Deep residual learning for image recognition, in: Proceedings of the IEEE Conference on Computer Vision and Pattern Recognition. pp. 770–778.

Jenkinson, M., Beckmann, C.F., Behrens, T.E., Woolrich, M.W., Smith, S.M., 2012. Fsl. Neuroimage 62, 782–790.

Ketkar, 2017. 12. Introduction to PyTorch - Deep Learning with Python: A Hands-on Introduction [Book] [WWW Document]. URL https://www.oreilly.com/library/view/deep-learning-with/9781484227664/A416804_1_En_12_Chapter.html (accessed 8.2.21).

Kim, H., Irimia, A., Hobel, S.M., Pogosyan, M., Tang, H., Petrosyan, P., Blanco, R.E.C., Duffy, B.A., Zhao, L., Crawford, K.L., Liew, S.-L., Clark, K., Law, M., Mukherjee, P., Manley, G.T., Van Horn, J.D., Toga, A.W., 2019. The LONI QC System: A Semi-Automated, Web-Based and Freely-Available Environment for the Comprehensive Quality Control of Neuroimaging Data. Front. Neuroinformatics 13, 60. https://doi.org/10.3389/fninf.2019.00060

Klein, S., Staring, M., Murphy, K., Viergever, M.A., Pluim, J.P., 2010. Elastix: a toolbox for intensity-based medical image registration. IEEE Trans. Med. Imaging 29, 196–205.

Küstner, T., Gatidis, S., Liebgott, A., Schwartz, M., Mauch, L., Martirosian, P., Schmidt, H., Schwenzer, N.F., Nikolaou, K., Bamberg, F., Yang, B., Schick, F., 2018. A machine-learning framework for automatic reference-free quality assessment in MRI. Magn. Reson. Imaging 53, 134–147. https://doi.org/10.1016/j.mri.2018.07.003

Manera, A.L., Dadar, M., Fonov, V., Collins, D.L., 2020. CerebrA, registration and manual label correction of Mindboggle-101 atlas for MNI-ICBM152 template. Sci. Data 7, 1–9.

Marek, K., Jennings, D., Lasch, S., Siderowf, A., Tanner, C., Simuni, T., Coffey, C., Kieburtz, K., Flagg, E., Chowdhury, S., Poewe, W., Mollenhauer, B., Klinik, P.-E., Sherer, T., Frasier, M., Meunier, C., Rudolph, A., Casaceli, C., Seibyl, J., Mendick, S., Schuff, N., Zhang, Y., Toga, A., Crawford, K., Ansbach, A., De Blasio, P., Piovella, M., Trojanowski, J., Shaw, L., Singleton, A., Hawkins, K., Eberling, J., Brooks, Deborah, Russell, D., Leary, L., Factor, S., Sommerfeld, B., Hogarth, P., Pighetti, E., Williams, K., Standaert, D., Guthrie, S., Hauser, R., Delgado, H., Jankovic, J., Hunter, C., Stern, M., Tran, B., Leverenz, J., Baca, M., Frank, S., Thomas, C.-A., Richard, I., Deeley, C., Rees, L., Sprenger, F., Lang, E., Shill, H., Obradov, S., Fernandez, H., Winters, A., Berg, D., Gauss, K., Galasko, D., Fontaine, D., Mari, Z., Gerstenhaber, M., Brooks, David, Malloy, S., Barone, P., Longo, K., Comery, T., Ravina, B., Grachev, I., Gallagher, K., Collins, M., Widnell, K.L., Ostrowizki, S., Fontoura, P., Ho, T., Luthman, J., Brug, M. van der, Reith, A.D., Taylor, P., 2011. The Parkinson Progression Marker Initiative (PPMI). Prog. Neurobiol., Biological Markers for Neurodegenerative Diseases 95, 629–635. https://doi.org/10.1016/j.pneurobio.2011.09.005

Mueller, S.G., Weiner, M.W., Thal, L.J., Petersen, R.C., Jack, C., Jagust, W., Trojanowski, J.Q., Toga, A.W., Beckett, L., 2005. The Alzheimer’s disease neuroimaging initiative. Neuroimaging Clin. N. Am. 15, 869–877.

Robin, X., Turck, N., Hainard, A., Tiberti, N., Lisacek, F., Sanchez, J.-C., Müller, M., 2011. pROC: an open-source package for R and S+ to analyze and compare ROC curves. BMC Bioinformatics 12, 1–8.

Russakovsky, O., Deng, J., Su, H., Krause, J., Satheesh, S., Ma, S., Huang, Z., Karpathy, A., Khosla, A., Bernstein, M., 2015. Imagenet large scale visual recognition challenge. Int. J. Comput. Vis. 115, 211–252.

Sørensen, T.J., 1948. A method of establishing groups of equal amplitude in plant sociology based on similarity of species content and its application to analyses of the vegetation on Danish commons. I kommission hos E. Munksgaard.

Sudlow, C., Gallacher, J., Allen, N., Beral, V., Burton, P., Danesh, J., Downey, P., Elliott, P., Green, J., Landray, M., 2015. UK biobank: an open access resource for identifying the causes of a wide range of complex diseases of middle and old age. PLoS Med. 12, e1001779.

Tremblay-Mercier, J., Madjar, C., Etienne, P., Poirier, J., Breitner, J., 2014. A PROGRAM OF PRE-SYMPTOMATIC EVALUATION OF EXPERIMENTAL OR NOVEL TREATMENTS FOR ALZHEIMER’S DISEASE (PREVENT-AD): DESIGN, METHODS, AND PERSPECTIVES. Alzheimers Dement. J. Alzheimers Assoc. 10, P808.

Van Essen, D.C., Ugurbil, K., Auerbach, E., Barch, D., Behrens, T.E.J., Bucholz, R., Chang, A., Chen, L., Corbetta, M., Curtiss, S.W., Della Penna, S., Feinberg, D., Glasser, M.F., Harel, N., Heath, A.C., Larson-Prior, L., Marcus, D., Michalareas, G., Moeller, S., Oostenveld, R., Petersen, S.E., Prior, F., Schlaggar, B.L., Smith, S.M., Snyder, A.Z., Xu, J., Yacoub, E., 2012. The Human Connectome Project: A data acquisition perspective. NeuroImage, ConnectivityConnectivity 62, 2222–2231. https://doi.org/10.1016/j.neuroimage.2012.02.018

Zijdenbos, A.P., Forghani, R., Evans, A.C., 2002. Automatic” pipeline” analysis of 3-D MRI data for clinical trials: application to multiple sclerosis. IEEE Trans. Med. Imaging 21, 1280–1291.

